# The impact of Subclinical Psychotic Symptoms on Delay and Effort discounting: insights from behavioral, computational, and electrophysiological methods

**DOI:** 10.1101/2023.06.24.546371

**Authors:** Damiano Terenzi, Massimo Silvetti, Giorgia Zoccolan, Raffaella I. Rumiati, Marilena Aiello

**Affiliations:** Institut de Neurosciences de la Timone, UMR 7289 CNRS & Aix-Marseille Université, Marseille, France; Computational and Translational Neuroscience Lab (CTNLab), Institute of Cognitive Sciences and Technologies, National Research Council (CNR), Rome, Italy; Area of Neuroscience, SISSA, Trieste, Italy; University of Rome Tor Vergata, Rome, Italy; Department of Psychology “Renzo Canestrari”, University of Bologna, Bologna, Italy

## Abstract

**Background:** The ability to value rewards is crucial for adaptive behavior and is influenced by the time and effort required to obtain them. Impairments in these computations have been observed in patients with schizophrenia and may be present in individuals with subclinical psychotic symptoms (PS).

**Methods:** In this study, we employed delay and effort-discounting tasks with food rewards in thirty-nine participants divided into high and low levels of PS. We investigated the underlying mechanisms of effort-discounting through computational modelling of dopamine prefrontal and subcortical circuits and the electrophysiological biomarker of both delay and effort-discounting alterations through resting-state frontal alpha asymmetry (FAA).

**Results:** Results revealed greater delay discounting in the High PS group compared to the Low PS group but no differences in the effort discounting task. However, in this task, the same levels of estimated dopamine release were associated with a lower willingness to exert effort for high-calorie food rewards in High PS participants compared to Low PS participants. Although there were no significant differences in FAA between the High PS and Low PS groups, FAA was significantly associated with the severity of participants’ negative symptoms.

**Conclusions:** Our study suggests that the dysfunction in temporal and effort cost computations, seen in patients with schizophrenia, may be present in individuals with subclinical PS. These findings provide valuable insight into the early vulnerability markers (behavioral, computational, and electrophysiological) for psychosis, which may aid in the development of preventive interventions.

## Introduction

When making decisions about rewards, considering the temporal and effort costs associated with obtaining the reward is crucial, as they can decrease its subjective value. Delayed rewards, for instance, require individuals to wait and forego immediate gratification, and can induce them to prefer immediate rewards even if smaller (Myerson and Green, 1995). At the same time, individuals may be less willing to engage in behaviors that require significant effort, despite the potential rewards being desirable (Botvinick *et al*., 2009). Delay discounting (DD) and effort discounting (ED) refer, respectively, to the devaluation of rewards that require more time or effort to be obtained and are mostly mediated by dopaminergic mesolimbic and frontal areas. In particular, the medial prefrontal cortex (mPFC), ventral striatum, posterior cingulate cortex and lateral parietal cortex correlate with delay discounting (Sellitto *et al*., 2010; Terenzi *et al*., 2021a, 2021b; Koban *et al*., 2023). Similarly, both cortical and subcortical regions are associated with effort discounting including motor areas, ventral striatum, anterior cingulate cortex, and anterior insula (Prevost *et al*., 2010; Lopez-Gamundi *et al*., 2021). Interestingly, a network of brain regions such as the mPFC, ventral striatum, and sensorimotor cortex have been observed to be commonly involved in processing rewards that require effort or are delayed (Aridan *et al*., 2019; Terenzi *et al*., 2022a).

Alterations in dopamine circuits and higher delay and effort discounting are common features of several disorders, including schizophrenia (SZ) (Amlung *et al*., 2019; Sellitto *et al*., 2022; Godefroy *et al*., 2023). Individuals with SZ show greater delay discounting rates (Brown *et al*., 2018; Amlung *et al*., 2019; Weinsztok *et al*., 2021). Similarly, they are even less willing to choose high-effort conditions as reward increases, and this behaviour is associated with negative symptom severity (Barch *et al*., 2014; Felice Reddy *et al*., 2016; Culbreth *et al*., 2018). Similar findings have been observed also in first-episode psychosis (Chang *et al*., 2020) as well as in individuals at clinically high risk for psychosis (Strauss *et al*., 2021). In our previous study, we found that altered delay and effort discounting could be observed even at earlier stages of the psychosis continuum, that is, in healthy individuals reporting subclinical psychotic experiences (Terenzi *et al*., 2019). These experiences also referred to as psychotic-like experiences (PLEs), are characterized by subthreshold forms of paranoid delusions, hallucinations, and negative symptoms and occur in healthy individuals from the general population without a clear psychotic disorder (Mossaheb *et al*., 2012; Terenzi *et al*., 2022b). Studies on the general adult population have indicated that approximately 8% of individuals reporting PLEs will develop psychotic symptoms within two years (Kelleher and Cannon, 2011), highlighting the potential significance of PLEs as a risk factor for developing psychotic disorders (Terenzi *et al*., 2019, 2022b).

Here, we investigate delay and effort discounting in individuals with subclinical psychotic symptoms by combining behavioural, computational, and electrophysiological methods. In particular, we use computational modelling to investigate the dynamics of effort discounting and dopamine transmission and examine the potential involvement of the mPFC in these processes. Computational modelling is a powerful tool that has been used to simulate the activity of the brain and investigate how it makes decisions, including in the context of schizophrenia (Rolls *et al*., 2008; Geana *et al*., 2022). Recently, Silvetti proposed a computational model focused on reward-based decision-making, specifically in the context of effort discounting of rewards (Silvetti *et al*., 2018, 2023). The model simulates the brain activity of the medial prefrontal cortex (mPFC) and the brainstem, and how noradrenaline and dopamine transmission influence behavior. It learns what actions lead to valuable outcomes (rewards) and exerts sufficient effort to carry out these actions successfully, with the mPFC playing a key role in learning when more effort is required via dopamine release. This model has been used to investigate individuals with autistic traits, providing insights into the underlying mechanisms of effort-based decision-making in these individuals (Goris *et al*., 2021) and suggesting the feasibility of the model to be applied also for other populations.

Moreover, in this study, we investigate the possible electrophysiological correlates of delay and effort discounting in subclinical psychosis using electroencephalography (EEG). Previous EEG studies have shown that greater left compared to right frontal alpha activity, known as left frontal asymmetry (LFA), is linked to motivated behavior (e.g., using an effort-discounting task) (Hughes *et al*., 2014) and trait impulsivity levels (Wendel *et al*., 2021). This index is also diminished in patients with SZ relative to healthy controls and correlates with the severity of negative symptoms (Jetha *et al*., 2009; Horan *et al*., 2014).

We hypothesized that individuals with high levels of subclinical psychotic symptoms (PS) (High PS group) would exhibit steeper delay and effort discounting compared to individuals with low PS (Low PS group). Using computational modelling, we hypothesized an altered underlying adjustment of effort allocation in the effort-discounting task as indexed by reduced estimated dopamine levels in High PS individuals relative to Low PS individuals. Finally, we expected to find reduced LFA in the High PS group compared to the Low PS group, and/or to observe a correlation between LFA and subclinical negative symptoms.

## Methods

### Participants

A sample of 368 participants was contacted through a social network and asked to fill out an internet-based version of the Community Assessment of Psychic Experiences (CAPE) (Konings *et al*., 2006). The CAPE is a 42-item questionnaire assessing self-reported subclinical positive, negative, and depressive symptoms in the general population (Mossaheb *et al*., 2012). Among this initial sample, 39 participants were selected based on the proposed cut-off score (using a score > 3.2 in the positive dimension) (Mossaheb *et al*., 2012; Terenzi *et al*., 2019). Specifically, 19 individuals with high subclinical positive symptoms (High PS participants), CAPE score > 3.2) and 20 with low subclinical positive symptoms (CAPE score < 2.8) were recruited. These two groups were matched for age, education, and BMI, as shown in Table 1. Participants were excluded based on specific criteria such as: (1) having a history of neurological or psychiatric illness; (2) having a family history of psychiatric illness in first-degree relatives; (3) using illicit substances (Modinos *et al*., 2010), (4) having eating disorders (assessed with the Eating disorder Inventory questionnaire, EDI-3) (Garner *et al*., 1983); and having food restrictions (e.g., vegetarianism, allergy, and others). All participants had a normal intelligence quotient measured through the Test di Intelligenza Breve (TIB) (Sartori *et al*., 1997) (mean: 104.93; SD: ±2.94; range: 98.50–111.29).

**Table 1.** Demographic and questionnaire data (mean and standard deviation). ^***^ = significantly different from Low PS, p < 0.001.

|  | High PS | Low PS |
| --- | --- | --- |
| **Demographics** |  |  |
| Age | 22.53 (2.01) | 22.70 (1.78) |
| Gender (female) | 17 | 16 |
| Education | 13.37 (1.26) | 13.90 (1.39) |
| BMI | 21.15 (2.23) | 22.56 (2.39) |
| **Psychotic-like experiences (CAPE)** |  |  |
| Positive symptoms | 3.55 (0.30)*** | 2.35 (0.19) |
| Negative symptoms | 4.30 (0.92)*** | 3.21 (0.63) |
| Depressive symptoms | 5.18 (0.89)*** | 3.80 (0.78) |
| Total Score | 13.03(1.75)*** | 9.36(1.49) |
| Bizarre experiences | 23.89 (4.83)*** | 14.90 (1.59) |
| Delusional ideation | 36.47 (3.24)*** | 24.15 (3.12) |
| Perceptual anomalies | 8.00 (2.52)*** | 6.00 (0.00) |
| **Cognition** |  |  |
| Digit Span Forward | 6.26 (0.87) | 6.45 (0.76) |
| **Mood/emotion** |  |  |
| BDI-II | 13.10 (6.98)*** | 6.85 (4.76) |
| BVAQ | 47.68 (8.53) | 47.75 (9.08) |
| **Impulsivity** |  |  |
| BIS-11 attentional | 18.47 (2.82)*** | 14.90 (2.99) |
| BIS-11 motor | 21.47 (3.29)*** | 18.60 (3.73) |
| BIS-11 non-planning | 24.63 (3.70) | 22.70 (4.41) |
| BIS-11 Total | 64.58 (7.10)*** | 56.20 (7.57) |
| **Reward sensitivity** |  |  |
| TEPS anticipatory | 39.53 (4.48) | 40.00 (3.64) |
| TEPS consummatory | 41.84 (4.95) | 43.95 (6.64) |

All participants gave written informed consent to participate in the study that was in line with the Declaration of Helsinki and approved by the SISSA Ethics Committee.

### Procedure

Before the test session, participants were requested to abstain from eating for at least two hours to induce comparable levels of hunger across all subjects. During the test session, we recorded participants’ self-reported hunger and fasting time and gathered their weight and height to compute their body mass index (BMI). We began the session by conducting a neuropsychological test to assess working memory, followed by several questionnaires that evaluated reward sensitivity, impulsivity, and mood. We then conducted a resting state EEG recording session before the participants performed, in a random order, one delay discounting task and one effort-based task.

### Food Delay-discounting Task

The food delay-discounting task requires making choices between immediate and delayed amounts of hypothetical food rewards. The task had 16 trials in total, with 4 choices per block, each block corresponding to a specific delay (1 hour, 2 hours, 5 hours, and 15 hours). The immediate option was adjusted to the participant’s discounting rate following a staircase procedure. For example, in every first trial of each block, the participant is presented with a choice between an immediate option of 6 units of food reward and a delayed option of 10 units of food reward after a certain delay. If the participant chooses the immediate option, the value of the immediate option is decreased in the next trial, making it less attractive. On the other hand, if the participant chooses the delayed option, the value of the immediate option is increased in the next trial, making it more attractive. This allows us to estimate the participant’s discounting rate. After four choices for a specific delay, the participant moved on to a new series of choices with another delay. The experimental procedure was similar to that used in a previous study (Terenzi *et al*., 2019). Participants were asked to select their preferred food from six popular snacks before starting the task. The purpose was to increase task ecological validity using snacks usually found in vending machines. One participant was unable to complete the task due to technical issues.

### Food Liking-rating task

Before the Effort-discounting task, participants were asked to taste and rate their liking of eight different real food items, comprising four high-calorie and four low-calorie foods, using an 11-point Likert scale. This task was used to choose the two preferred foods for each participant - one high-calorie and one low-calorie - which were subsequently presented as visual stimuli in the Effort-discounting task as pictures of the real foods (Terenzi *et al*., 2019).

### Effort-discounting task

In this task, participants had to earn points to obtain their two favourite foods by pressing a left or right key on the keyboard corresponding to the left or right part of the screen displaying the food. Next, they had to maintain a constant contraction of force through a hand-grip dynamometer for 14 seconds after each decision to earn 1 point/gram of the selected food.

Importantly, the amount of force required to get the high-calorie food increases during the task while the one related to the low-calorie food was kept constant. The task consisted of three schedules of 15 choices each, meaning that a total of 45 points/grams of food had to be earned and eaten at the end of the task. In the first schedule (*easy schedule*), the proportion of force to keep constant to get 1 point/gram of both food options was equal to 8% of the participant’s maximal voluntary contraction (MVC). In the following two schedules (*medium and hard schedules*), the amount of effort required for the fruit/vegetable option was kept at 8% while the force associated with the choice of the snack option increased (13% and 33% in the medium and hard schedule, respectively) (see Figure 1). Real-time feedback on the exerted force was provided for each trial. For a similar procedure see (Giesen *et al*., 2010; Chong *et al*., 2017) The effort discounting task of the current study was designed to remove any temporal confounds by keeping the same duration (14 seconds) in all effort trials across the three effort conditions.

**Figure 1.**
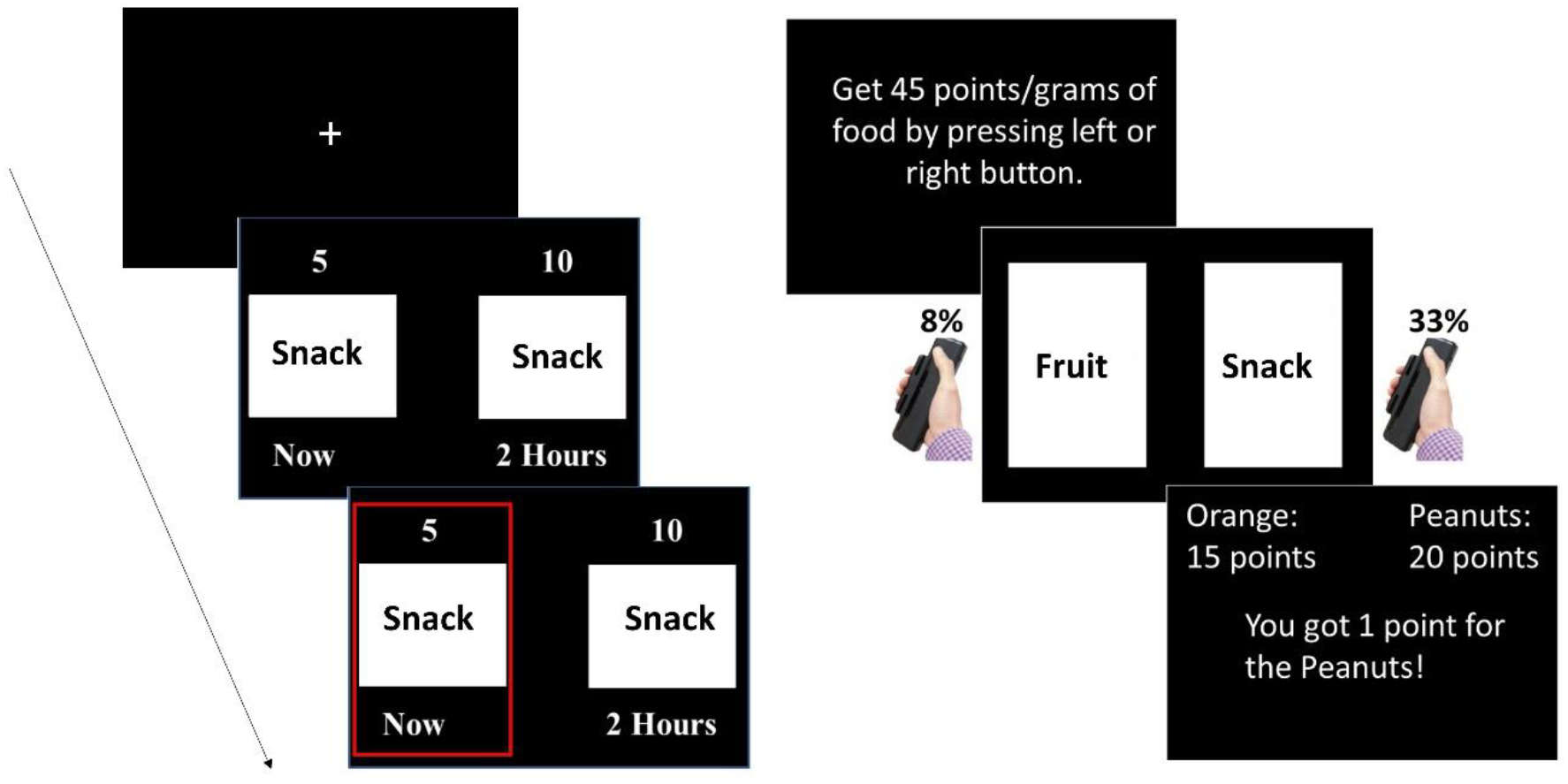
(Left) Delay-Discounting task. The task involved presenting subjects with a choice between receiving a small reward immediately or a larger reward after a delay, with a fixation period of 1 second before each trial. After making their choice, the preferred option remained highlighted for 1 second. (Right) Example trial sequence of the hard schedule of the Effort-discounting task. The amount of force for low-calorie items stays at 8% of participants’ MVC, whereas the % of force for snacks increased during the task (from 8% in the easy schedule to 33% in the hard schedule). During the task, participants aimed to obtain a total of 45 points of food. They could choose to allocate these points to both options or allocate them solely to one option.

### Computational modelling: the Reinforcement Meta Learner (RML)

The equations governing the RML and a detailed description can be found in Silvetti et al. (2023) and in the Supplementary Materials (Silvetti *et al*., 2023). We used the RML to simulate human behaviour during the Effort-discounting task (Figure 2). As the native architecture of the model does not represent time discounting, we simulated only the Effort discounting task. For each choice, the RML was asked to choose between a high reward/high effort option (snack) and a low reward/low effort option (fruit). The probability of selecting the high-effort option during one trial depended on the control signal (boost) selected by the MPFC_Boost_ during that trial, and the boost level depended on the trade-off between the expected reward and the intrinsic cost of boosting. Large boost values determine large NE and DA values that, in turn, promote effortful actions and task engagement (Figure 2). We set the reward magnitude equal to 2 for snacks and 1 for fruit, and the motor cost (hand grip squeezing) was set to .5 (for simulating 8% MVC), 1 (13% MVC), and 2 (33% MVC). The overall task structure and the role of reward and motor cost in the equations determining action selection are described in detail in Simulation 2 of Silvetti et al. (Silvetti *et al*., 2018).

**Figure 2.**
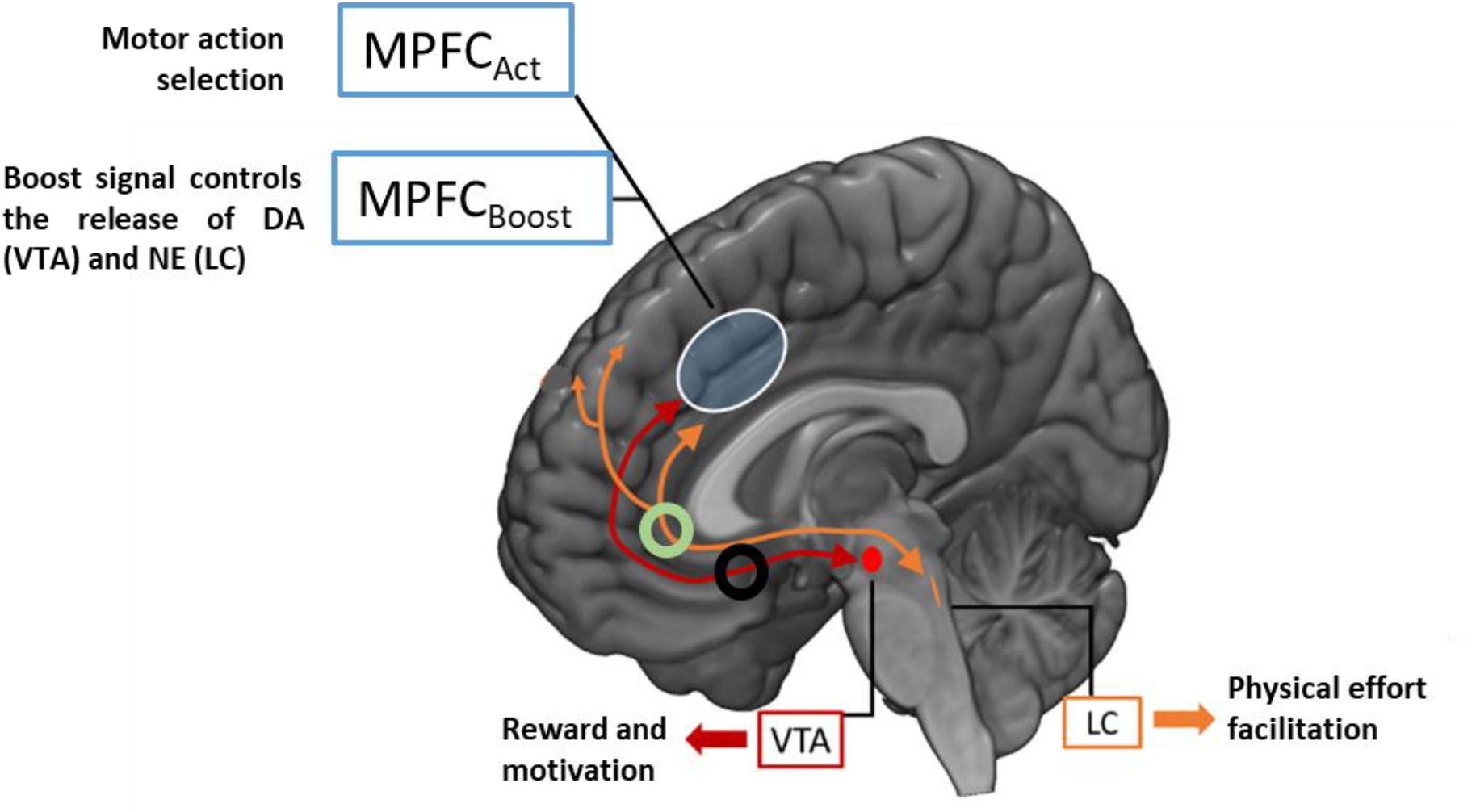
Functions and neuroanatomical mapping of RML modules. The MPFC_Act_ selects motor actions (e.g. «snack» option selection and grip squeezing) based on state-action values. The MPFC_Boost_ exploits the same machinery, but it selects instead the boost signal targeting the VTA and the LC, and modulating their output. Trial-by-trial update of MPFC modules’ belief on state-action values is performed via Bayesian learning. *DA*^*M*^ (black disk) and *NE*^*M*^ (green disk) are two free parameters multiplying the output of VTA and LC respectively. We optimized these two parameters for each human participant by fitting the RML behavioural responses with those of the participant.

We fitted the RML behaviour on the behaviour of each participant during the Effort-discounting task, generating personalized simulations of both neural and behavioural patterns (digital twins). To this aim, we optimized two RML parameters (*DA*^*M*^ and *NE*^*M*^) representing the activity levels of VTA and LC (Figure 2). These parameters were two real numbers multiplying the output of VTA and LC modules, so that if they were greater than 1 they amplified the output, while they decreased it if smaller than 1. In this way, we could estimate the strength of both DA and NE effects on decision-making for each participant. Parameters optimization was conducted by minimizing, for each participant, the mean squared error (MMSE) between the percentage of “snack” option selection by RML for each schedule *s* (*RML_sk*_*s*_) with that by human (*Human_sk*_*s*_). The objective can be written as:

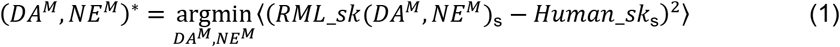

Where *RML*_*sk*(*DA*^*M*^, *NE*^*M*^)_s_ is the percentage of “snack” option selected by the RML, during the schedule *s*, and given the pair *DA*^*M*^ and *NE*^*M*^. Finally, ⟨… ⟩ indicates mean, while * indicates optimality. Before the start of each estimation process, the RML was trained to learn the value of the two options (fruit and snack).

### EEG data acquisition and analysis

See Supplementary Materials.

### Questionnaires and neuropsychological tests

See Table 1 and Supplementary Materials.

### Statistical analyses

Data were analyzed in R (https://www.r-project.org/). The Shapiro-Wilk test was used to confirm normal distribution. Independent samples T-tests or Mann-Whitney U tests were used for group comparisons on demographic, neuropsychological, and questionnaire measures, while the chi-square test was used for gender comparisons. A linear mixed-effects model was used to predict discounting rates. Furthermore, we calculated the area under the empirical discounting curve (AUC) as a model-free parameter (Myerson *et al*., 2001) for each participant and entered these values in a repeated measures ANOVA. Liking rating scores were compared using repeated measures ANOVA, and a Generalized Linear Mixed-Effects model (GLMM) with a logistic link function was used to predict the probability of accepting or rejecting to exert effort for high-calorie foods across the different effort schedules (easy, medium, hard) and groups (high PS vs low PS). All models included participants’ ID as a random intercept (see Supplementary Materials). EEG alpha asymmetry scores and power spectra were compared across groups using linear mixed models, and correlations were performed to examine relationships between demographic, questionnaire, neuropsychological, behavioral, and EEG measures.

The Reinforcement Meta Learner (RML) was used to estimate the trade-off between the value and cost of effort and its link with neuromodulators such as dopamine (Silvetti *et al*., 2018, 2023) and the activity of the medial prefrontal cortex (see above). Correlations between CAPE negative and depressive scores and our measures of impulsivity, motivation, and mood were performed considering all participants together. However, when performing correlations related to the CAPE-Positive subscale, separate analyses were performed within each group as the two groups were explicitly selected based on predetermined cutoff criteria for the CAPE-positive symptom subscale, which resulted in the identification of individuals with either extremely high or extremely low scores on this subscale.

## Results

### Demographic and questionnaire data

The two groups of participants were matched for gender, age, education, and BMI (*ps* > 0.065). They did not differ in subjective ratings of hunger (*p* > 0.162). As expected, participants with High PS scored significantly higher than participants with low PS on all the CAPE sub-scales (all *ps* < 0.001) and the BDI-II scale (*p* = 0.002). No differences emerged in the Digit Span forward test (*p* = 0.479), on the TEPS anticipatory (*p* = 0.270) and consummatory scales (*p* = 0.718), as well as on the BVAQ questionnaire (*p* = 0.981). Lastly, High PS participants showed higher levels of impulsivity on the BIS-11 questionnaire compared to Low PS participants (*p* = 0.001) (See Table 1). See supplementary information for detailed statistics.

### Food Delay-Discounting task

The linear mixed model showed a main effect of time delays (χ^2^(3) = 17.563, *p* < 0.001) and a significant interaction between time delays and group (χ^2^(3) = 14.594, *p* = 0.002) (Figure 3). Post hoc tests showed that participants’ subjective values for food rewards were statistically different for delays that were more distant in time: from 1 hour to 5 hours (t(519) = 3.542, *p* = 0.002) and from 1 hour to 15 hours (t(519) = 3.467, *p* =0.002); while no differences emerged between the other subjective values across the time delays (*Ps* > 0.068). Post-hoc analysis on the triple interaction showed a higher discounting rate for food rewards in High PS compared to Low PS for a medium-high time delay of 5 h (t(55.9) = -2.691, *p* = 0.0437)

**Figure 3.**
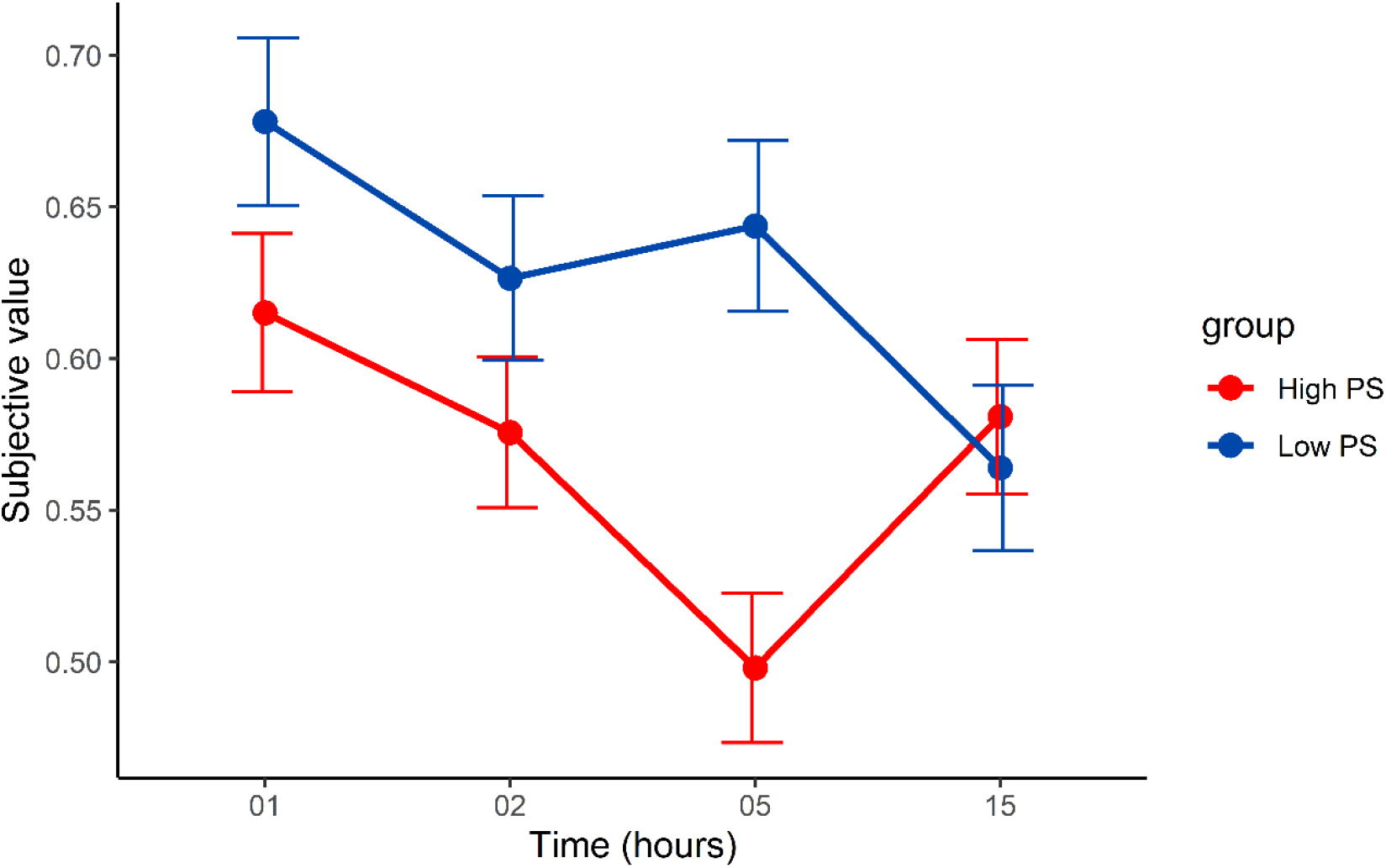
Delay discounting function for subjective values across time delays (in hours) for High PS (in red), and Low PS (in blue). The x-axis represents the time delays in hours, and the y-axis represents the subjective values for food rewards. The error bars represent SEM.

### Food liking rating task

A repeated measures ANOVA examining the effects of group (high PS, low PS) and calorie content (high-calorie, low-calorie) on liking ratings did not yield any significant results (all *Ps* > 0.525).

### Effort-Discounting Task

A first generalized linear mixed-effects logistic regression (glmer) model showed a main effect of the Effort Schedules (χ^2^(2) = 8.891, p = 0.012). Post-hoc tests showed that participants were more willing to accept to exert effort for high-calorie food in the medium effort schedule compared to the easy (β = 0.541, *p* < 0.001) and hard (β = 0.319, *p* = 0.037) effort schedules. No differences between the easy and hard schedules emerged (β = -0.222, *p* = 0.117). A second glmer model including a further predictor such as the estimated levels of DA for each participant in each effort schedule of the task showed a main effect of the levels of DA (χ^2^(1) = 24.003, *p* < 0.001) indicating that higher levels of DA predicted a higher acceptance to exert effort for high-calorie foods across the effort schedules and groups. Importantly, this model also showed a significant triple interaction Group ^*^ Effort Schedules ^*^ levels of DA (χ ^2^ (2), *p* < 0.001). Post-hoc analysis revealed that higher levels of DA in High PS participants were associated with a lower acceptance of effort for reward compared to Low PS participants (β = -1.322, *p* = 0.002) in the hardest effort schedule of the task (Figure 4). Notably, adding the levels of DA improves model fit as shown by Akaike information criterion (AIC of the first model = 1871.779; AIC of the second model = 1792.561). A further glmer model including the estimated levels of norepinephrine for each participant in each effort schedule did not show significant differences between the two groups of participants. Lastly, repeated measures ANOVA on the estimated activity of the mPFC across the different effort schedules and groups did not reveal any significant results.

**Figure 4.**
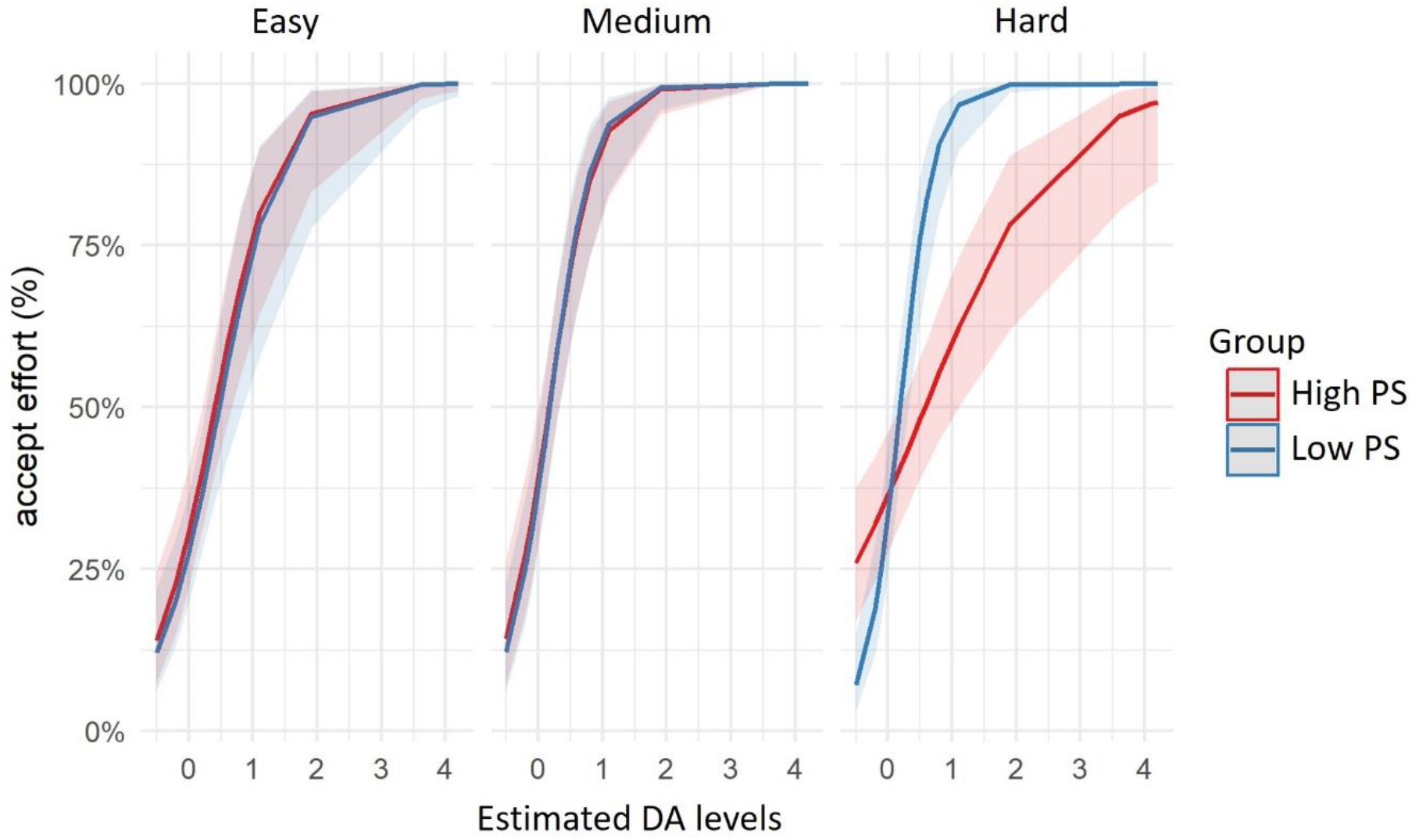
Probability of accepting to exert effort (%) for high-calorie food, as a function of the estimated dopamine (DA) levels, across three different effort schedules: easy, medium, and hard. The red line corresponds to the High PS group, while the blue line corresponds to the Low PS group.

### Resting state EEG

A linear mixed model was performed on alpha asymmetry values computed from frontal electrodes as well as from control electrodes (medial and parietal), with the factors Group (High PS and Low PS) and session (eyes closed and open). Results did not show any significant effects (*ps* > 0.089). Similarly, a linear mixed model on the delta, theta, alpha, and beta relative power frequencies across the sessions (eyes closed and open) did not show a main effect of the group nor a group per band frequency interaction (*ps* > 0.293). Importantly, the CAPE questionnaire’s score of negative symptoms, a self-reported index of altered motivation, positively correlated with our electrophysiological index of motivation such as the left frontal asymmetry (rho = 0.37, p = 0.023; Figure S1) (low alpha recorded during the eyes closed session).

### Correlational analyses

We found significant correlations between the subscales of the CAPE and the experimental tasks. Bizarre experience scores were positively correlated with increased motivation for high-calorie foods, while delusional ideation scores were associated with greater impulsivity. Furthermore, higher mPFC estimated activity during the effort task was linked to higher depressive symptoms and lower alexithymia scores (see Supplementary Materials for detailed statistics).

## Discussion

The purpose of this study was threefold. First, to replicate our prior research demonstrating altered delay and effort discounting of rewards in individuals with subclinical psychosis. Second, to apply computational modelling of dopamine prefrontal and subcortical circuits to deepen our understanding of effort-discounting of rewards in subclinical psychosis. Third, to investigate the potential of resting-state alpha asymmetry as an electrophysiological biomarker of delay and effort discounting alterations in this population. By incorporating behavioral, computational, and electrophysiological measures, this comprehensive approach enhances the understanding of the phenomena at study.

In line with our hypothesis, we observed that High PS group exhibited a greater discounting rate for food rewards compared to the Low PS group in particular at a medium-high time delay of 5 hours. This result is consistent with previous research showing a steeper delay discounting of rewards in patients with SZ relative to healthy controls (Brown *et al*., 2018; Purcell *et al*., 2022). Our results also suggest that using an ecological task involving food rewards (popular snacks) and short time delays (hours instead of days or months) can be a useful tool to study delay discounting. This task may be particularly advantageous because it involves more tangible and immediate choices, and it may better capture the real-world decision-making context of individuals’ food choices. However, our results suggest that the effects of delay discounting may differ depending on the delay interval. This result offers valuable insights into determining the optimal time for discounting rewards in individuals with subclinical psychotic disorders.

In examining effort computations, we found that the two groups of participants did not show differences in their willingness to exert effort for high-calorie food rewards across the different effort schedules. At variance with the previous study (Terenzi *et al*., 2019), the task designed for this study used fewer grams of food rewards to get and therefore fewer number trials (45 vs. 100) which may have limited the task’s sensitivity in detecting any potential differences between the two groups. Interestingly, when simulating the trade-off between the value and effort cost encoded by the mesolimbic/mesocortical dopamine network through computational modelling, we found that higher estimated dopamine release predicted a higher acceptance to exert effort for high-calorie foods and that this effect differed between individuals with high versus low levels of subclinical psychotic symptoms. Indeed, the same levels of dopamine release were associated with a lower willingness to exert effort for high-calorie food rewards in High PS participants compared to Low PS participants. Importantly, this difference was observed only in the high-effort condition and not in the other conditions, suggesting that only when the effort required is high, individuals with High PS may show an inefficient dopamine function. This finding is consistent with previous research suggesting that SZ is associated with dopamine dysregulation and reduced motivation and reward processing (Barch et al., 2014). According to Michely and colleagues, dopamine is important for integrating physical effort with reward value, which helps individuals to maximize rewards (Michely *et al*., 2020). Studies using fMRI on healthy individuals have suggested that a dopamine circuit including the ventral striatum and mPFC is involved in effort discounting (Botvinick *et al*., 2009; Aridan *et al*., 2019), and a PET study linked dopamine function in these regions to the willingness to exert effort (Treadway *et al*., 2012). In SZ, previous fMRI studies examining effort discounting have consistently found abnormal function in the ventral striatum and prefrontal cortex (Huang *et al*., 2016; Culbreth *et al*., 2020; Prettyman *et al*., 2021). Notably, dysfunctional activation in frontostriatal brain regions during the monetary incentive delay task (MID) has also been observed in adolescents with subclinical psychotic symptoms (Papanastasiou *et al*., 2018). Overall, our study suggests that alterations in the mesolimbic/mesocortical dopamine network may serve as a potential biomarker for vulnerability to psychosis.

To our knowledge, our study is the first to investigate also frontal alpha asymmetry as a biomarker of motivational impairment in individuals with subclinical psychotic symptoms. Contrary to our hypothesis, we did not find any differences in frontal alpha asymmetry between High PS and Low PS groups. While some studies have reported altered FAA in patients with SZ compared to controls (Jetha *et al*., 2009; Horan *et al*., 2014; Jang *et al*., 2020) a recent study on youth at clinically high risk for psychosis did not find such differences (Bartolomeo *et al*., 2019). Nonetheless, this study has found a relationship between FAA and the severity of negative symptoms. Similarly, we observed that higher levels of resting FAA (indicating less alpha power in the left frontal hemisphere compared to the right) were associated with greater severity of negative symptoms. This finding suggests, first, that the relationship between FAA and negative symptoms is not due to illness chronicity, as our participants were healthy individuals reporting high levels of subclinical psychotic symptoms. Secondly, as our participants were healthy individuals, they were antipsychotic-naïve. Thus, the previously observed association between altered FAA and negative symptoms in SZ may not be solely attributable to medication effects.

Finally, our study also uncovered intriguing correlations between different facets of subclinical psychosis, such as bizarre experiences and delusional ideation, and measures of motivation and impulsivity. Individuals with higher levels of subclinical psychotic symptoms and more pronounced bizarre experiences exhibit a heightened motivation to obtain high-calorie foods, as evidenced by their increased willingness to work for such rewards in the effort discounting task. Conversely, individuals with higher levels of delusional ideation exhibit greater impulsivity, as reflected in their reduced ability to delay gratification in the delay discounting task. These correlations underscore the complex interplay between subclinical psychosis and motivation and impulsivity, shedding light on potential avenues for further research and therapeutic intervention. We also observed a positive correlation between the participants’ estimated activity of the mPFC during the effort task and their depressive symptoms, as well as a negative correlation with alexithymia scores. The mPFC is important for decision-making and emotion regulation (Dixon *et al*., 2017) and abnormal function has been linked to depression ((Pizzagalli and Roberts, 2022) and alexithymia (Riadh *et al*., 2019). The effort discounting task that we employed may be useful for assessing this relationship, and further research on mPFC activity during such tasks could provide insights into affective abnormalities related to impaired decision-making and emotion regulation.

Some limitations to our study should be noted. First, we did not correct for multiple comparisons in our correlational analyses, which could increase the likelihood of Type I errors. Therefore, future studies with larger sample sizes are needed to replicate and confirm our findings. Second, it would be informative to use an effort-discounting task with a wider range of reward amounts to better understand the relationship between effort and reward in individuals with subclinical psychotic symptoms.

In conclusion, our study indicates that subclinical psychotic symptoms can affect decision-making, particularly in the temporal and effort discounting of rewards. Dysregulation of the frontostriatal dopamine network may be involved in the motivational deficits observed in psychosis. Further research is needed to confirm the predictions derived from our computational model. Lastly, the electrophysiological marker we identified, known as FAA, provides insight into the neural mechanisms underlying negative symptoms in subclinical psychosis. These findings highlight the importance of examining behavioral, computational, and electrophysiological measures to better understand the complex nature of subclinical psychotic symptoms and their impact on decision-making. Overall, our study contributes to the growing body of literature on the early detection and prevention of psychotic illness and may have potential implications for the development of targeted interventions for individuals at risk of developing psychosis.

