## Supplementary materials for "The impact of Subclinical Psychotic Symptoms on Delay and Effort discounting: insights from behavioral, computational, and electrophysiological methods"

### Statistical analyses

Data were analyzed in R (https://www.r-project.org/). The Shapiro-Wilk test was used to confirm normal distribution. Independent samples T-tests or Mann-Whitney U tests were used for group comparisons on demographic, neuropsychological, and questionnaire measures, while the chi-square test was used for gender comparisons. A linear mixed-effects model was used to predict discounting rates using the lmer function in the lm4 package (Bates et al., 2015). P-values were obtained from the analysis of variance type 3 tests (ANOVA). Predictors were one inter-subject (the two groups: high PS vs low PS), and one intra-subject (the four different time delays: 1 h, 2, h, 5h, 15 h). Furthermore, we calculated the area under the empirical discounting curve (AUC) as a model-free parameter (Myerson et al., 2001) for each participant and entered these values in a repeated measures ANOVA.
Liking rating scores were compared using repeated measures ANOVA, and a Generalized Linear Mixed-Effects model (GLMM) with a logistic link function was used to predict the probability of accepting or rejecting to exert effort for high-calorie foods across the different effort schedules (easy, medium, hard) and groups (high PS vs low PS). All models included participants' ID as a random intercept and post-hoc comparisons were performed using emmeans (estimated marginal means) package in R. EEG alpha asymmetry scores and power spectra were compared across groups using linear mixed models, and correlations were performed to examine relationships between demographic, questionnaire, neuropsychological, behavioral, and EEG measures.
The Reinforcement Meta Learner (RML) was used to estimate the trade-off between the value and cost of effort and its link with neuromodulators such as dopamine (Silvetti et al., 2018, 2023) and the activity of the medial prefrontal cortex (see above). Correlations between CAPE negative and depressive scores and our measures of impulsivity, motivation, and mood were performed considering all participants together. However, when performing correlations related to the CAPE-Positive subscale, separate analyses were performed within each group (rather than analyzing all participants together) as the two groups were explicitly selected based on predetermined cutoff

criteria for the CAPE-positive symptom subscale, which resulted in the identification of individuals with either extremely high or extremely low scores on this subscale.

#### EEG data acquisition and analysis

We used an EEG procedure to measure resting-state FAA as an index of approach motivation following a method commonly used in previous studies (Allen et al., 2004; Harmon-Jones et al., 2018; Metzen et al., 2022). EEG signals were recorded using a BioSemi ActiveTwo system with 64 channels at a sampling rate of 1024Hz and preprocessed offline with EEGLAB v 2021.1 (Delorme and Makeig, 2004), implemented in Matlab R2021n. The alpha power spectrum (8-13Hz) was calculated using a Fast Fourier Transformation, and the alpha asymmetry index was obtained by measuring the difference of log-transformed alpha between homologous left and right electrodes. Frontal (F4 & F3), medial (C4 & C3), and parietal (P4 & P3) electrodes were included in the analysis. Delta, theta, alpha, and beta relative power frequencies were also extracted for additional exploratory analyses (Fuggetta et al., 2014; Howells et al., 2018). Two 3.5-minute sessions with eyes closed and eyes open were administered in a counterbalanced order. Resting state EEG was not recorded in two subjects due to technical problems.

#### Questionnaires and neuropsychological tests

Participants completed the following questionnaires: the Temporal Experience of Pleasure Scale (TEPS) (Gard et al., 2007), the Beck Depression Inventory (Beck et al., 1996), the Barrat Impulsiveness Scale (BIS-11) (Fossati et al., 2011). Furthermore, they were asked to perform the Digit Span Forward test (Orsini et al., 1987) to assess their working memory. Lastly, participants performed the Bermond-Vorst Alexithymia Questionnaire (BVAQ) (Bermord et al., 1994) since it has been shown that higher scores on alexithymia are associated with a higher risk of developing psychosis (van der Velde et al., 2015).

#### Computational modelling: the Reinforcement Meta Learner (RML)

The RML (Silvetti et al., 2018,2023) is an autonomous agent that operates within a meta-RL (Doya 2002) and Bayesian (Kalman 1965) frameworks and comprises four modules forming a system deputed to reward and information-based decision-making. In previous works, we have shown that the RML is capable of adaptive behaviour and near-optimal decision-making in many tasks leveraging on a wide range of different cognitive processes, like visual attention, working memory, effort-based decision-making, and non-stationary foraging (Silvetti et al., 2018). Moreover, the RML has been shown effective in predicting both behavioural and neural dynamics of human and nonhuman primates (Silvetti 2018, 2019, 2020a, 2020b, 2023). The equations governing the RML and a detailed description can be found in Silvetti et al. (2023), in this work we did not implement any modification to the model. The RML architecture is inspired by the neurophysiology of a cortical-subcortical circuit including the medial prefrontal cortex (MPFC) and the catecholamine nuclei (Figure 2). Its architecture consists of four computational modules, two modules simulate subcortical structures – respectively, the ventral tegmental area (VTA) for midbrain DA signals conveying primary and non-primary reward-related information, and NE signals from the locus coeruleus (LC) that modulate cognitive and physical effort. Two additional modules simulate the roles of the MPFC in generating, respectively, motor actions directed to the external environment ($MPFC_{Act}$) and "executive" actions that regulate the internal environment by boosting brainstem neuromodulators NE and DA ($MPFC_{Boos}$t). The $MPFC_{Boost}$ module selects the optimal level of cognitive control and/or physical effort using iterative updates of state-action values, and it comes with an intrinsic cost, which increases with the intensity of the boost signal. For this reason, the RML objective is to maximize reward while minimizing the boost cost.

$$(DA^M, NE^M)^* = \underset{DA^M, NE^M}{\operatorname{argmin}} \langle (RML\_sk(DA^M, NE^M)_{\mathrm{s}} - Human\_sk_{\mathrm{s}})^2 \rangle \tag{1}$$

Where $RML\_sk(DA^M, NE^M)_{\mathrm{s}}$ is the percentage of "snack" option selected by the RML, during the schedule *s*, and given the pair $DA^M$ and $NE^M$. Finally, $\langle \ldots \rangle$ indicates mean, while * indicates optimality. Before the start of each estimation process, the RML was trained to learn the value of the two options (fruit and snack).

**Demographic and questionnaire data**

The High PS group was not significantly different from the Low PS group in terms of age ($t(37) = -0.280$, $p = 0.781$), gender ($\chi^2(1) = 0.671$, $p = 0.412$), education ($t(37) = -1.100$, $p = 0.279$), BMI (t(37)

= -1.902, p = 0.065), and self-reported hunger levels(t(37) = 1.426, p = 0.162). Participants with High PS scored significantly higher than Low PS participants in CAPE positive t(37) = 15.026, p < .001), CAPE negative (t(37) = 4.338, p = < 0.001), CAPE depressive (t(37) = 5.156, p < 0.001) scores. Furthermore, High PS participants scored higher than Low PS participants in all CAPE positive subscales such as Bizarre experiences (t(37) = 7.898, p < .001), Delusional ideation (t(37) = 12.111, p < .001), and Perceptual anomalies (t(37) = 3.557, p < 0.001). The High PS also showed higher BDI scores compared to the Low PS group (t(37) = 3.289, p = 0.002). No differences emerged between the High PS and Low PS groups on the Digit Span forward test (t(37) = -0.715, p = 0.479), the TEPS anticipatory scale (t(37) = -1.119, p = 0.270), TEPS consummatory scale (t(37) = -0.363, p = 0.718), and the BVAQ questionnaire (t(37) = -0.023, p = 0.982). However, High PS participants demonstrated significantly higher levels of impulsivity on the BIS-11 questionnaire compared to Low PS participants (t(37) = 3.289, p = 0.001).

**Correlational analyses**

Several correlations were found in the study between measures of impulsivity, motivation, and mood, with psychotic-like experiences.

First, the delusional ideation subscale of the CAPE questionnaire showed a negative correlation with AUC values in the delay-discounting task among High PS individuals (rho = -0.54, p = 0.021; Figure S1A), indicating that decisional impulsivity was higher in High PS participants with higher levels of delusional ideation. This correlation was not observed among Low PS participants (rho = -0.00, p = 0.757). In addition, High PS participants' bizarre experiences positively correlated with their effort allocation to obtain rewards in the hardest schedule of the effort task (rho = 0.48, p = 0.039) (Figure S1B). This correlation was only found to be significant in the High PS group, and not in the Low PS group (*ps* > 0.260).

When analyzing CAPE depressive scores among all participants we found a positive correlation with the estimated mPFC activity in the hardest effort schedule (rho = 0.33, p = 0.042). Conversely, the estimated mPFC activity in the hardest effort schedule showed a negative correlation with BVAQ scores (rho = -0.32, p = 0.044).

Lastly, the CAPE questionnaire's score of negative symptoms among all participants, a self-reported index of altered motivation, positively correlated with our electrophysiological index of motivation such as the left frontal asymmetry (rho = 0.37, p = 0.023; Figure S1C) (low alpha recorded during the eyes closed session).

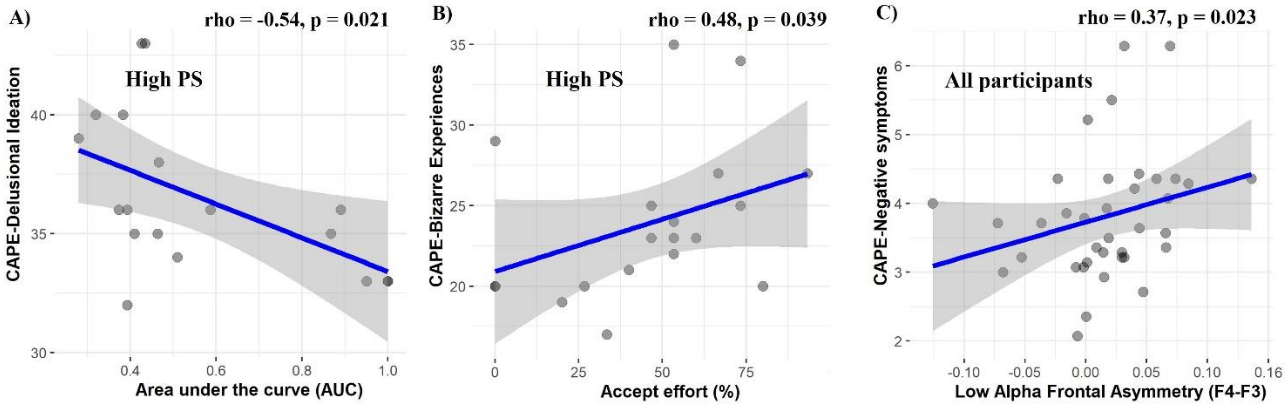


**Figure S1. Correlations.** A) Cape-Delusional ideation scores of High PS participants negatively correlated with their AUC scores in the delay-discounting task (rho =-0.54, p = 0.021). B) Cape-Bizarre experiences scores of High PS participants positively correlated with their willingness to exert effort in the effort-discounting task (rho = 0.48, p = 0.039). C) Cape-Negative symptoms scores across participants positively correlated with their low Alpha Frontal Asymmetry (rho = 0.37, p = 0.023).
